# Cancer-Associated Mutations Enhance The Sensitivity Of The Trupath Gα_Q/11_ System

**DOI:** 10.1101/2022.09.01.506210

**Authors:** Dewi Safitri, Matthew Harris, Abigail Pearce, Xianglin Huang, Matthew Rosa, Kerry Barkan, Edward Wills, Maria Marti-Solano, Matthew D. Falk, Graham Ladds

**Affiliations:** Department of Pharmacology, University of Cambridge, Tennis Court Road, Cambridge CB2 1PD, United Kingdom; Pharmacology and Clinical Pharmacy Research Group, Institut Teknologi Bandung, Bandung 40132, Indonesia; Takeda Development Centre Americas Inc, San Diego, CA, USA

**Keywords:** GPCRs, class B, G protein dissociation, calcium, G_q/11_, GIPR, GLP-1R, OXTR

## Abstract

G protein-coupled receptors (GPCRs) are the largest family of cell surface receptors and are a common drug target. They can be stabilised in different conformational states by ligands to activate multiple transducers and effectors leading to a variety of cellular responses. The potential of agonists to activate select pathways has important implications for drug discovery. Thus, there is a clear need to profile the initial GPCR signal transduction event, activation of G proteins, to enhance understanding of receptor coupling and guide drug design. The BRET-based biosensor suite, TRUPATH, was recently developed to enable quantification of the activation profiles of all non-visual G proteins (excluding G_olf_ and G_14_) and has since been utilised in numerous studies. However, it fails to detect G_q/11_ activation for a number of GPCRs previously reported to display promiscuous secondary coupling to G_q/11_. Here we report modifications to the Gα_q_ and Gα_11_ biosensors in the switch I region that prevent intrinsic GTPase activity (R183C/Q). Except for the PAC1R, substitution with cancer-associated mutations, Cys or Gln, significantly increased sensitivity to allow detection of robust, reliable, and representative G_q/11_ responses to Class B1 GPCRs. We also demonstrate the utility of these modified biosensors for promiscuously coupled class A GPCR that have primary G_s_-coupling. Thus, we propose that modification to Gα_q/11_ may also be necessary in other biosensor systems to enable detection of G_q/11_ activation.

## INTRODUCTION

G protein-coupled receptors (GPCRs) represent the largest family of plasma membrane proteins and are important therapeutic targets (Hauser, Avet, Normand, Mancini, & Inoue, 2022; Katritch, Cherezov, & Stevens, 2013; Santos et al., 2016). Many GPCRs are known to signal pleiotropically, whereby activation results in the stimulation of multiple signalling pathways due to promiscuous G protein activation (Avet et al., 2022; Hauser et al., 2022; Inoue et al., 2019). The potential of agonists to stabilise particular receptor conformations that functionally bias a GPCR to a particular signalling pathway (biased agonism) has important implications for drug discovery and highlights the need to understand this receptor pleiotropy (Smith, Lefkowitz, & Rajagopal, 2018). Measurement of downstream signalling events (e.g., cAMP accumulation, intracellular calcium release (Ca^2+^)_I_ or phosphorylation of effector proteins such as extracellular signal-regulated kinase/mitogen-activated protein kinase (ERK/MAPK), p38 or protein kinase B (Akt)) has long been used to experimentally evaluate agonist induced GPCR activation (Goldsmith & Dhanasekaran, 2007; Wootten, Miller, Koole, Christopoulos, & Sexton, 2017). Interpretation of results can, however, be complicated by accumulation of second messengers and crosstalk between pathways. Therefore, measurement of the initial signal transduction event, G protein activation, can provide important additional information to illuminate the mechanisms underlying GPCR signalling. One of the first methods for measuring G protein activation utilised a radiolabelled, non-hydrolysable version of guanine nucleotide triphosphate (GTP), called [^35^S]GTPɣS (Asano, Katada, Gilman, & Ross, 1984; Hilf, Gierschik, & Jakobs, 1989) which would accumulate following exchange for guanine nucleotide diphosphate (GDP) at activated G proteins (Gilman, 1987; Harrison & Traynor, 2003). However, [^35^S]GTPɣS assays are commonly performed using membrane preparations and have the hazards associated with radioactivity.

Resonance energy transfer (RET) techniques have become increasingly popular as tools to probe almost every step of GPCR signalling pathways in cells, including interaction between receptor and ligands, G proteins, or effectors; β-arrestin recruitment; second messenger production; receptor dimerization; receptor translocation; and activation of transcription factors (Audet et al., 2008; Ayoub et al., 2007; Harikumar, Happs, & Miller, 2008; Inoue et al., 2019; Jiang et al., 2007; Jimenez-Vargas et al., 2018; M. J. Lohse et al., 2008; Martin J. Lohse, Nuber, & Hoffmann, 2012; Olsen et al., 2020; Picard, Schönegge, Lohse, & Bouvier, 2018; Surdo et al., 2017). Not only do RET-based assays allow the move away from radioactivity and offer the advantage to monitor protein-protein interaction in living cells, but they also enable quantification of kinetic aspects of GPCR signalling.

Bioluminescence RET (BRET) was first utilised to assess heterotrimeric G protein activation by Galès et al., by measuring BRET between receptor and G protein (Galés et al., 2005). Since then, numerous luminescence-based assays utilising different methods have been developed to measure G protein activation, including miniG protein, NanoBiT, enhanced bystander BRET, BERKY, G-CASE, and TRUPATH (Audet et al., 2008; Avet et al., 2022; Galés et al., 2005; Inoue et al., 2012, 2019; Maziarz et al., 2020; Nehmea et al., 2017; Nobles, Benians, & Tinker, 2005; Olsen et al., 2020; Philip, Sengupta, & Scarlata, 2007; Schihada, Shekhani, & Schulte, 2021). The latter, TRUPATH system, exploits the dissociation of Gα-GTP from Gβɣ to measure G protein activation. Optimized sensors consisting of Gα-Rluc8 and Gβɣ-GFP2 were developed for all non-visual G proteins, with the exception of G_olf_ and G_14_, and included the first sensors for G_15_ and G_gust_ (Olsen et al., 2020). TRUPATH has since been employed to assess the G protein activation profile of a number of GPCRs (Knight et al., 2021; Lieb et al., 2021; Singh, Senatorov, Cheshmehkani, Karmokar, & Moniri, 2022; Voss, Mahardhika, Inoue, & Müller, 2022; Wall et al., 2022).

Pleiotropic signalling has been documented for numerous GPCRs (Avet et al., 2022; Hauser et al., 2022; Knight et al., 2021; Lieb et al., 2021; Olsen et al., 2020; Singh et al., 2022; Voss et al., 2022; Wall et al., 2022). Class B1 GPCRs, for example, are classically considered Gα_s-_coupled GPCRs but have now been demonstrated to consistently exhibit secondary G protein coupling (Wootten et al., 2017). Indeed, many class B1 GPCRs have been shown to stimulate typical Gα_q/11_-mediated signalling events, such as intracellular Ca^2+^ mobilisation and IP1 production (Clark et al., 2021; Drissi, Oise Lasmoles, Ronique Le Mellay, Marie, & Le Lieberherr, 1998; Harris et al., 2021; Hinkle, Donnelly, Cody, Sheldon, & Isfort, 2005; Putney & Tomita, 2012; Wootten et al., 2017). However, for two of these receptors, gastric inhibitory polypeptide receptor (GIPR) and glucagon like peptide-1 receptor (GLP-1R), despite evidence that intracellular Ca^2+^ mobilisation is Gα_q/11_-dependent, no Gα_q_ coupling was detected in two recent studies (Avet et al., 2022; Jones et al., 2021)

Here, we investigate Gα_q/11_ activation by GIPR and GLP-1R using the TRUPATH system and identify that activation can only be detected upon mutation of a critical residue in the switch I region of Gα_q/11_ (R183), which results in impaired G protein inactivation. Through screening all remaining class B1 receptors with Cys and Gln mutated Gα_q/11_, we report the optimal modifications to TRUPATH to enable detection of G protein activation for each receptor. Furthermore, we demonstrate that mutation of Gα_q/11_ may also have applications for measuring G_q/11_ activation of class A GPCRs that are primarily coupled to G_s_. Importantly, this study also highlights the potential for analogous modifications to other G protein activation biosensor systems.

## MATERIAL AND METHODS

### DNA Constructs

c-myc-β_3_-AR was a gift from Aylin Hanyaloglu (Imperial College London). β_2_-AR was a gift from Asuka Inoue (Tohoku University), whereas A_2A_R was from David Poyner (Aston University), D_3_R and μ-OR were purchased from cDNA resource centre (cdna.org). SmBiT-VPACR1 was generated via PCR amplification of VPACR1 sequence, without the native signal peptide, which was ligated into a pcDNA3.1(+)-sigSmBiT backbone, containing the signal peptide of the murine 5-HT_3_ receptor. The remaining receptor constructs and FLAG-tagged RAMPs were used as previously described (Harris et al., 2021). Plasmid pNL[NFAT-RE-NlucP-Hygro] was purchased from Promega (UK) to determine NFAT-mediated transcription by Gα_q/11_-coupled receptors. To assess Gα_q/11_ coupling to GPCRs, TRUPATH BRET biosensors were purchased from Addgene. This kit contains plasmids encoding Gα-Rluc8, Gβ, and Gɣ-GFP2 subunits of heterotrimeric G protein complex.

### Peptides and compounds

The following human peptides: GLP-2 (1-33), CRF, PTH (1-34), GHRH, VIP, calcitonin, αCGRP, AM and AM2 were purchased from Bachem (Bubendorf, Switzerland). Human GIP (1-42) was provided by Takeda Pharmaceuticals (San Diego, US). Human glucagon and GLP-1 (7–36)NH_2_ were custom synthesised by Generon (UK). Secretin was purchased from Cambridge Bioscience (UK). PACAP-27 was obtained from Selleckchem (UK). All peptides were prepared as 1 mM stocks in water containing 0.1% BSA. Oxytocin and isoprenaline were purchased from Sigma (UK), dopamine hydrochloride was purchased from Tocris (UK) and these compounds were prepared as 10 mM stocks in water containing 0.1% BSA. [D-Ala^2^, *N*-MePhe4, Gly-ol]-enkephalin (DAMGO), purchased from Sigma, and CGS21680, purchased from Tocris (UK) were prepared as 10 mM stocks in DMSO. YM-254890 was purchased from Wako Chemicals Inc (USA) and was dissolved in DMSO at 10 mM stock.

### Cell Culture

Human embryonic kidney (HEK) 293T cells were cultured at 37°C in a 5% CO_2_ humidified incubator in Dulbecco’s modified Eagle’s medium (DMEM)/F12 (1:1) with GlutaMAX™ (Thermo Fisher, UK) media supplemented with 10% (v/v) foetal bovine serum (FBS, Sigma, UK), and 1% antibiotic antimycotic (Sigma, UK). Cells were routinely passaged before reaching confluency. HEK293T cells stably expressing pNL[NFAT-RE-NlucP-Hygro] were also maintained in a complete DMEM/F12 medium supplemented with 0.06 mg/ml hygromycin (Thermofisher, UK). HEK293T cells overexpressing full-length GIPR or GLP-1R with an N-terminal FLAG tag were purchased from Multispan, Inc (Hayward, CA) and cultured as per the manufacturer’s protocol in DMEM with 10% FBS (GE Healthcare, Pittsburgh, PA) and 1 µg/mL puromycin (ThermoFisher, Waltham, MA). Cells were transfected using polyethyleneimine (PEI, Polysciences, Germany) and 150 mM NaCl using a 1:6 (w:v) ratio of DNA:PEI, unless otherwise stated.

### Site-directed mutagenesis

pcDNA5/FRT/TO-GAlphaQ/11-RLuc8 plasmid (part of “TRUPATH” G protein dissociation assay kit) served as the template for making Gα_q_ and Gα_11_ mutants. Single point mutagenesis was performed using QuikChange Lightning site-directed mutagenesis kit (Agilent Technologies LDA UK Limited, UK) in accordance with the manufacturer’s instruction. The oligonucleotides for mutagenesis were designed using the manufacturers online primer design tool (https://www.agilent.com/store/primerDesignProgram.jsp)

Three different mutants for Gα_q_ were created using mutagenic primers *forward* 5’-CTGTGGTGGGGACGCAAACTCTAAGCACATCTTGTTGCGT-3’ and *reverse* 5’-ACGCAACAAGATGTGCTTAGAGTTTGCGTCCCCACCACAG-3’ to generate R183C; *forward* 5’-CTGTGGTGGGGACTTTAACTCTAAGCACATCTTGTTGCGT-3’ and *reverse* 5’-ACGCAACAAGATGTGCTTAGAGTTAAAGTCCCCACCACAG-3’ to generate R183K, and *forward* 5’-CCCTGTGGTGGGGACTTGAACTCTAAGCACATC-3’ and *reverse* 5’-GATGTGCTTAGAGTTCAAGTCCCCACCACAGGG-3’ to generate R183Q. Whereas the following primers were used to generate Gα_11_ R183C: *forward* 5’-GGTGGGCAC GCAGACCCGCAGCA-3’ and *reverse* 5’-TGCTGCGGGTCTGCGTGCCCACC-3’, R183K: *forward* 5’-GCCGGTGGTGGGCACTTTGACCCGCAGCACGTC-3’ and *reverse* 5’-GACG TGCTGCGGGTCAAAGTGCCCACCACCGGC-3’, and R183Q variants: *forward* 5’-CGGTGGTGGGCACTTGGACCCGCAGCAC-3’ and *reverse* 5’-GTGCTGCGGGTCCAAGTGCCCACCACCG-3’, respectively.

### Measurement of intracellular calcium (Ca^2+^)_i_ mobilisation

HEK293T cells were seeded at a density of 200,000 cells/well on black 96-well plates (Greiner Bioscience, UK) coated with poly-L-lysine (PLL, Caltag Medsystem, UK). Transfection was performed using Fugene HD (Promega, UK) according to the manufacturer’s instructions, using a 1:3 (w:v) ratio of DNA:FuGene. Cells were then assayed for (Ca^2+^)_i_ release in response to ligands in the range of 0.1 nM to 10 μM as previously described (Harris *et al*., 2021). Data were normalized to the maximum intensity produced by 10 μM ionomycin (Cayman Biosciences, UK). To confirm that (Ca^2+^)_i_ responses were G_q/11_-mediated, cells were pretreated for 30 mins in 100 nM of YM-254890, which inhibits G_q/11_ signalling (Takasaki *et al*., 2004).

### IP-One Accumulation Assay

IP1 response was measured using the Cisbio IP-One Gq HTRF kit (Bedford, MA), according to the manufacturer’s protocol. Briefly, HEK 293T-FLAG-GIPR/-FLAG-GLP-1R cells or HEK293T cells transiently transfected with OXTR were washed two times in 1x stimulation buffer and resuspended to a concentration of 2.86 × 10^6^ cells/mL (GIPR response), 7.14 × 10^6^ cells/mL (GLP-1R response) or 2.9 × 10^6^ cells/mL (OXTR response) in 1x stimulation buffer. 7 µL of cell suspension was dispensed in low volume, white, 384-well polypropylene plates (Corning, Tewksbury, MA), followed by addition of 7 µL peptide agonist diluted in 1x stimulation buffer to the final indicated concentration. Cells were stimulated for 2 hr (GIPR and OXTR response) or 3 hr (GLP-1R response) prior to addition of 3 µL IP1-d2 and 3 µL anti-IP1 Tb cryptate antibody diluted in lysis buffer. Plates were read on a BMG Labtech PHERAstar plate reader (Ortenberg, Germany), with excitation at 337 nm and emission at 615 nm and 665 nm.

### TRUPATH G protein dissociation assay

HEK293T cells were seeded at the density of 10^6^ cells/well in a 6-well plate and were grown overnight in complete DMEM/F12 1:1 Glutamax ™ (Thermofisher, UK). On the next day, cells were transfected with native or mutant Gα_q_-Rluc8, Gβ_3_, Gɣ_9_, the receptor of interest, and pcDNA3.1 at a ratio of 1:1:1:1:1 with a total of 2 μg DNA per well. For Gα_11_, Gɣ_13_ was used in place of Gɣ_9_ (Olsen *et al*., 2020). For CLR, pcDNA3.1 was replaced with a N-terminally FLAG-tagged RAMP. After 24 hours, cells were trypsinised and re-seeded onto poly-L-lysine (Sigma, UK)-coated white 96-well plates (Greiner, UK) at a density of 50,000 cells/well and cultured overnight. On the day of assay, cells were washed with HBSS before addition of 80 μl HBSS, 20 mM HEPES, and 0.1% BSA, adjusted to pH 7.4. 10 μl coelenterazine 400a (Nanolight Technology, USA) was added to a final concentration of 5 μM and the plate was incubated in the dark for 5 minutes to reach the equilibrium. 10 μl of ligand (0.01 nM to 1 μM) was then added and the BRET signal recorded for 20 minutes on a Mithras LB940 multimode plate reader allowing sequential integration of signal detected from GFP2 and Rluc8. The BRET ratio corresponds to the ratio of light emission from GFP2 (515 nm) over Rluc8 (400 nm). The BRET ratio was corrected to vehicle and the net BRET ratio at the 11-minute time-point used to produce the dose-response curves shown. To verify that Gα_q/11_ mutants were still sensitive to YM-254890, experiments were performed where cells were pretreated with 100 nM YM254890.

### Nuclear factor of activated T cells (NFAT) reporter assay

HEK293T-NFAT cells were transfected with 2 µg of plasmid expressing either OXTR, GIPR, or GLP-1R using Lipofectamine 2000 for 6.5 hours (Thermofisher, UK), according to the manufacturer’s instructions, using a 1:2 (w:v) ratio of DNA:Lipofectamine 2000. Media and transfection reagent were then replaced with the complete media. 30-hour post-transfection, cells were plated onto a PLL-coated white 96-well plate at a density of 200,000 cells/well in complete DMEM/F12. Approximately 48 hours after transfection, cells were stimulated for 6.5 hours with ligands in a range of 0.1 pM to 1 µM in DMEM/F12 media containing 0.5% FBS at 37 °C with 5% CO_2_. After stimulation, media was removed and replaced with 50 µL per well 0.01 mM Furimazine or NanoGlo® Lytic Buffer (diluted 1 in 500, Promega, UK). After a 10-minute shaking incubation protected from light, unfiltered luminescence signal was recorded using a CLARIOstar® plate reader (BMG, UK) using 0.5 second interval reading time.

### Data Analysis

Data analyses for all experiments was performed using GraphPad Prism version 9.0.0 (GraphPad Software, San Diego, California USA). All dose-response data were fitted using the non-linear three parameter logistic equation to obtain values for pEC_50_ and E_max_. All experiments were performed at least three times in duplicate unless otherwise indicated. Data are expressed as mean ± standard error of the mean (SEM) and statistical differences were determined, at p<0.05, using a one-way ANOVA followed by Dunnett’s post-hoc test or Students’ t-test, as appropriate.

## RESULTS

### GIPR, GLP-1R, and OXTR stimulate Gα_q/11_-mediated signalling events

Class B1 GPCRs are classically considered G_s_-coupled receptors, whereby activation leads to the production of cAMP. However, there is now substantial evidence that class B1 GPCRs have promiscuous G protein couplings and significant pleiotropic signalling (Alexander et al., 2019; Avet et al., 2022; Hauser, Attwood, Rask-Andersen, Schiöth, & Gloriam, 2017; Pándy-Szekeres et al., 2018). For example, activation of a number of class B1 GPCRs has been demonstrated to result in the mobilisation of (Ca^2+^)_i_ from intracellular stores and that this is, at least in part, G_q/11_-dependent (Clark et al., 2021; Drissi et al., 1998; Harris et al., 2021; Hinkle et al., 2005; Wootten et al., 2017). Indeed, for two typical class B1 GPCRs, GIPR and GLP-1R, we detect a dose-dependent IP1 production, as well as increase in (Ca^2+^)_i_ mobilisation and NFAT-dependent gene expression in response to stimulation with their respective cognate agonists (GIP (1-42) and GLP-1 (7-36)NH_2_)) (Figure 1A-C). NFAT was reported to be an inducible transcription factor activated through G_q_ pathway (Boss, Talpade, & Murphy, 1996). Stimulation of cells expressing the primarily G_q/11_-coupled GPCR, oxytocin receptor (OXTR) with oxytocin, similarly resulted in a dose-dependent increase in IP1, (Ca^2+^)_i_, and NFAT-dependent gene expression. Pre-treatment of cells with YM-254890, a cyclic depsipeptide that selectively inhibits Gα_q/11_ activation by preventing the exchange of GDP for GTP (Takasaki et al., 2004), resulted in almost complete attenuation of GIP (1-42), GLP-1 (7-36)NH_2_ and oxytocin-mediated (Ca^2+^)_i_ mobilisation (Figure 1B). This, therefore, indicates that GIPR and GLP-1R are, like OXTR, G_q/11_-coupled GPCRs.

**Figure 1.**
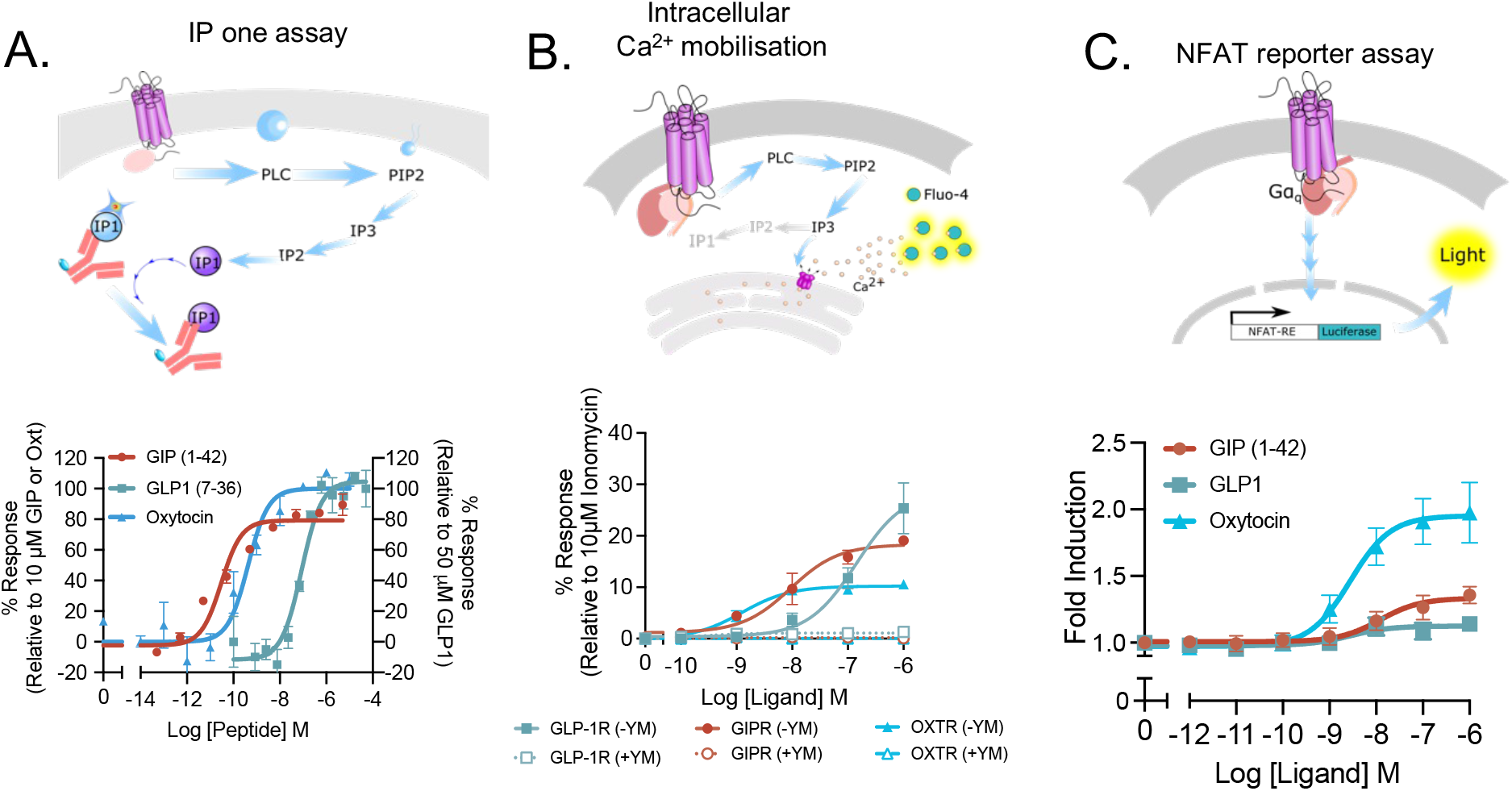
GIPR, GLP-1R, and OXTR activation dose-dependently promotes Gq-mediated downstream signalling. (**A**-**C**) Schematics demonstrating the principles of the IP one assay (**A**), (Ca^2+^)_i_ mobilisation detection using Fluo-4/AM (**B**) and the NFAT reporter assay (**C**). (**D**) HEK293T cells transiently transfected with GIPR, GLP-1R or OXTR were stimulated with GIP (1-42), GLP-1 (7-36)NH_2_ or oxytocin, respectively, and IP one production (**D**), (Ca^2+^)_i_ mobilisation (**E**) or NFAT-dependent gene expression (**F**) measured. Data are the mean ± SEM from at least 3 individual repeats. IP one responses were normalised to the response to the highest concentration of ligand as 100%. (Ca^2+^)_i_ mobilisation release from the endoplasmic reticulum (ER) was measured in the presence and absence of YM-254890 and data expressed relative to 10 μM ionomycin. NFAT-dependent gene expression was measured as the fold induction of luciferase expression.

We next set out to measure G_q/11_ protein activation at GIPR and GLP-1R using the recently developed, open-source G protein BRET biosensor suite, TRUPATH (Olsen et al., 2020) (illustrated in Figure 2A). As expected, stimulation of OXTR with oxytocin triggered a dose-dependent loss of BRET for G_q_ and G_11_, bought about by Gα-Rluc8 and Gβɣ-GFP2 dissociation upon G protein activation (Figure 2C and D, native). However, contrary to expectation, there was no significant change in the net BRET ratio for GIPR or GLP-1R, even when stimulated with a maximal concentration of agonist (1 μM).

**Figure 2.**
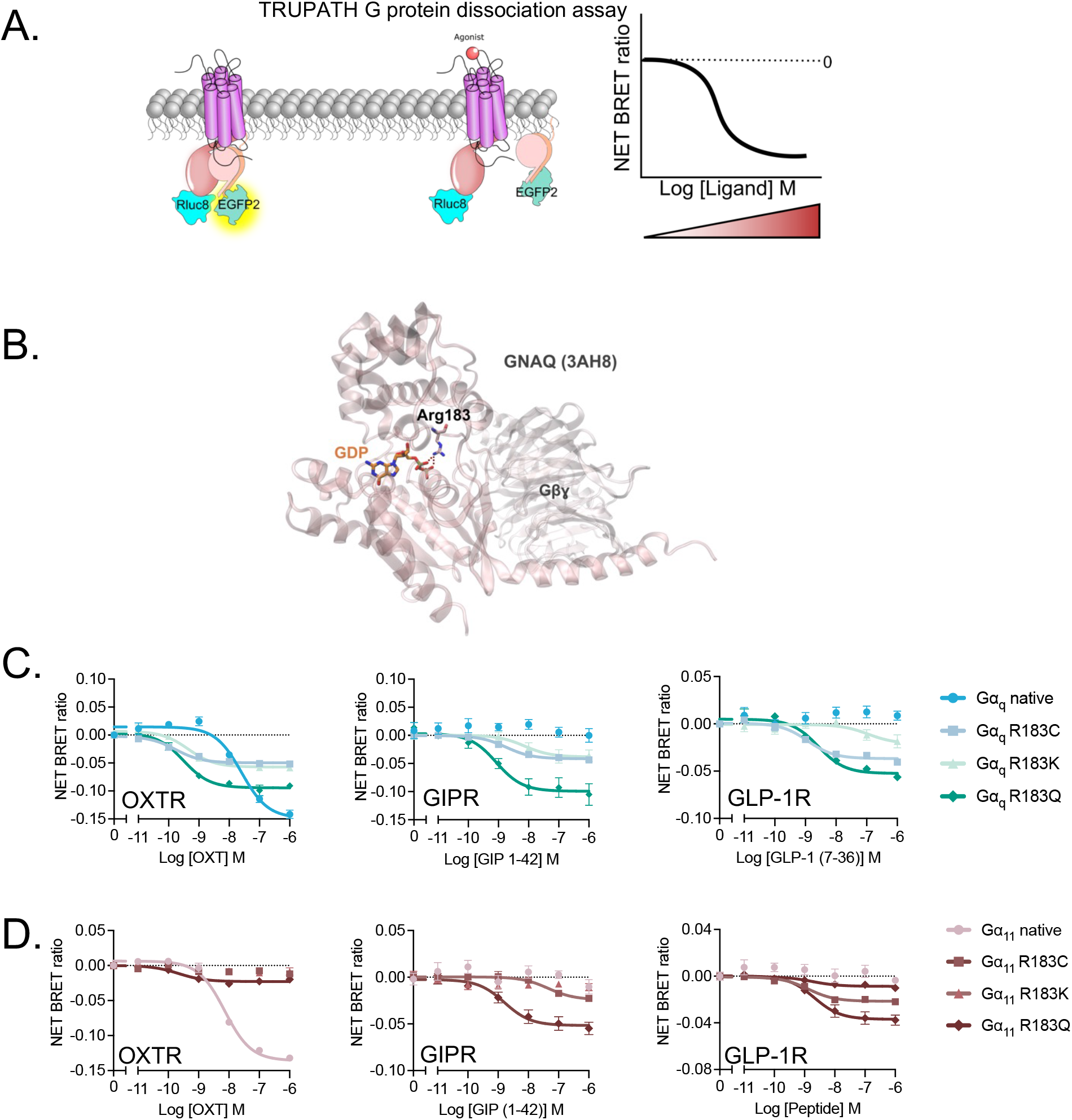
Mutation to R183 in the switch I region enables detection of Gα_q/11_ activation at GIPR and GLP-1R. (**A**) Schematic illustration of TRUPATH G protein dissociation assay (Olsen 2020). (**B**) A structure of inactive G_q_ protein heterotrimer highlighting the direct interaction between the Arg residue and GDP. (**C**-**D**) G protein dissociation in HEK 293T cells transiently expressing native, R183C, R183Q and R183K G_q_ (**B**) and G_11_ (**C**) TRUPATH biosensors and OXTR, GIPR or GLP-1R in response to GIP (1-42), GLP-1 (7-36)NH_2_ or oxytocin. Data are the mean ± SEM of 3-6 individual repeats performed in duplicate.

### Mutation of R183 in Gα_q/11_ allows detection of GIPR and GLP-1R-mediated G_q/11_ activation

Codon 183 in exon 4 of *GNAQ* and *GNA11* is located in the GTP binding region of Gα_q/11_ and plays an important role in the hydrolysis of GTP to GDP to inactivate the G protein. Several point mutations have been found to exist at this position that result in reduced or ablated GTP hydrolysis and increased activity of G_q/11_. Due to the proposed role of G_q/11_ as an oncogene (Parish et al., 2018), it is, thus, not surprising that mutations to R183 have been observed in many cancers and have been found to induce spontaneous tumour metastasis in mouse models (Böhm et al., 2013; Murali, Wiesner, Rosenblum, & Bastian, 2012; O’Hayre et al., 2013; Raamsdonk et al., 2010; Thomas et al., 2016; Van Raamsdonk et al., 2009). Two of the most common point mutations are p.183R>C in *GNA11* and p.183R>Q in *GNAQ*. Given the enhanced downstream signalling detected for these mutants (Bastin et al., 2012; Schrage et al., 2015; Thomas et al., 2016), we hypothesised that mutating R183 to prevent GTP hydrolysis would slow Gα and Gβɣ reassociation to increase the dynamic range of the TRUPATH assay for weakly coupled GPCRs. The arginine residue is also found to be a direct interaction site for GDP and is conserved across Gα subunits as depicted in Figure 2B. We, therefore, mutated R183 to cysteine or glutamine for both G_q_ and G_11_. Furthermore, as a control to verify the importance of a positive charge at position 183 in stabilising the transition state of GTP hydrolysis, we also generated the R183K mutation.

For G_q_ (Figure 2C), mutation of R183 to cysteine enabled detection of a dose-dependent reduction in net BRET for GIPR and GLP-1R, although the range was still small in comparison to other Gα biosensors. At GIPR and GLP-1R, the glutamine mutant displayed activation with ~1.2 and ~2-fold greater E_max_, respectively (p<0.001), but similar potency to R183C, indicating that R183Q is the optimal biosensor. The Lys mutation displayed a similar maximal level of activity to Cys but with slightly reduced potency for GIPR, whilst for GLP-1R the potency and E_max_ were significantly reduced (p<0.01). This indicates that a large proportion of the GTPase activity is retained when the positively charged Lys is present. Interestingly, for OXTR, the maximal response was reduced but oxytocin potency increased for all three mutants compared to the native G_q_.

To validate that the G protein dissociation observed for GIPR and GLP-1R was not an artifact, we pre-treated with YM-254890. Indeed, this abolished both GIPR and GLP-1R-mediated activation of Gα_q_-mutants and demonstrated that substitution at position 183 does not impact sensitivity to YM-254890 (Supplementary figure S1).

For G_11_ (Figure 2C), the GIPR response displayed a similar, ~2-fold greater E_max_ for the Gln mutant compared to the Cys mutant, whilst the Lys mutant was not significantly different to native G_11_. G_11_ activation at GLP-1R was also ~2-fold greater for the Gln mutant compared to the Cys mutant although in this case there was only very weak activity with the Lys mutant. Again, mutation of Arg to Cys, Gln or Lys in G_11_ attenuated the oxytocin induced reduction in BRET ratio at the OXTR, although in this case there was an almost complete loss of signal. This is possibly due to the different location of the Rluc8 insertion in Gα_11_ compared to Gα_q_. Overall, this indicates that it is likely that the cancer-related mutations in Gα_q/11_ increase the sensitivity of the system but reduce the total activatable pool of G proteins. The reduced availability of associated heterotrimeric G proteins is evidenced by the observation that the raw BRET ratio is significantly lower in Q and C substituted G_q/11_ compared to native (Supplementary figure S2). Together, this enables detection of G_q/11_ responses for GPCRs that primarily couple to and activate other G proteins and also weakly activate G_q/11_ but reduces the maximal signalling window for GPCRs that couple primarily to G_q/11_.

To ensure that the mutant G_q_ and G_11_ biosensors did not promote unselective receptor coupling thus giving a false readout, we verified each mutant against the β_3_-adrenoceptor (β_3_AR), dopamine D_3_ receptor (D_3_R), and μ-opioid receptor (μ-OR), all of which are reported not to couple to G_q/11_ (gpcrdb.com). Upon stimulation of β_3_AR, D_3_R or μ-OR there was no significant reduction in the net BRET ratio for native G_q/11_ or any of the amino acid substitutions at position 183 (Supplementary figure S3-5). There was, however, the expected reduction in net BRET ratio for G_s_ (for β_3_AR), G_oA_ (for D_3_R) or G_i1_ (for μ-OR) upon stimulation with isoprenaline, dopamine or DAMGO, respectively. These data demonstrate that all receptors are functional and do not couple with native or mutant Gα_q_ or Gα_11_. Thus, whilst these mutant G_q/11_ proteins are known to inactivate to a lower extent than their wildtype counterparts they can only be activated by receptors that endogenously couple to G_q/11_.

### Differential Gα_q/11_ coupling to class B1 GPCRs

Having established the potential of mutating position R183 in Gα_q_ and Gα_11_ to allow detection of activation by GIPR or GLP-1R, we next sought to investigate G_q/11_ activation at all class B1 GPCRs. Here, we screened native, R183C and R183Q G_q/11_ activation for the remaining 13 class B1 GPCRs as well as the complex between CLR and 3 different RAMPs (CLR:RAMP1; CGRP receptor (CGRPR), CLR:RAMP2; adrenomedullin receptor (AMR), and CLR:RAMP3; adrenomedullin2/intermedin receptor (AMR2). This allowed us to systematically monitor class B1 activation of both G_q_ (Figure 3) and G_11_ (Figure 4). The potency (pEC_50_) and the range of BRET reduction or span are summarised in Figure 5.

**Figure 3.**
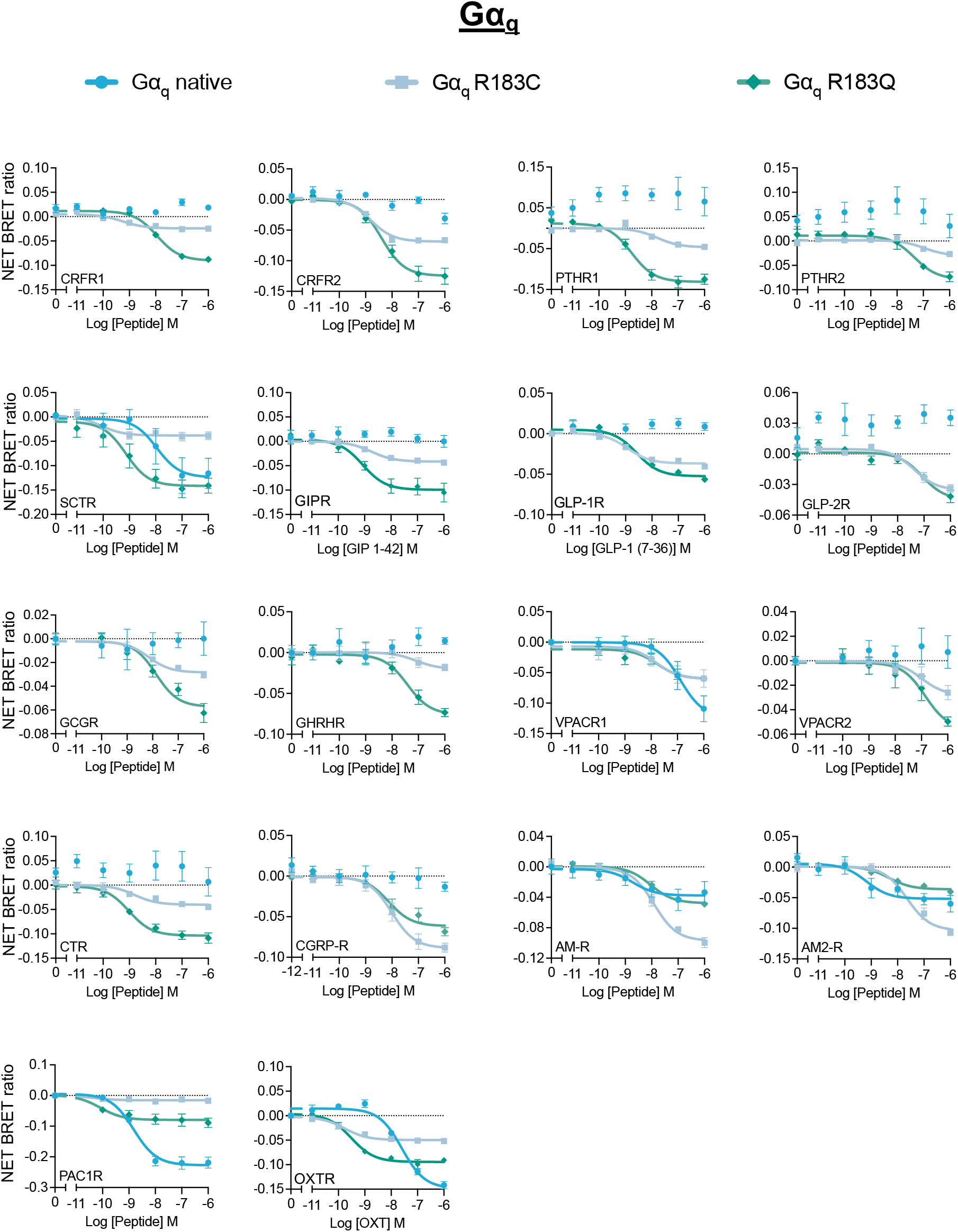
Differential responses of all class B1 GPCRs using G_q_ native and mutants in TRUPATH assay. Dose-response curves of dissociation of native or mutant Gq for all class B1 GPCRs. HEK 293T cells transiently transfected with each class B1 GPCR and native, R183C or R183Q G_q_ TRUPATH biosensors were stimulated with their cognate agonist. Data are the mean ± SEM of 3-6 individual repeats performed in duplicate.

**Figure 4.**
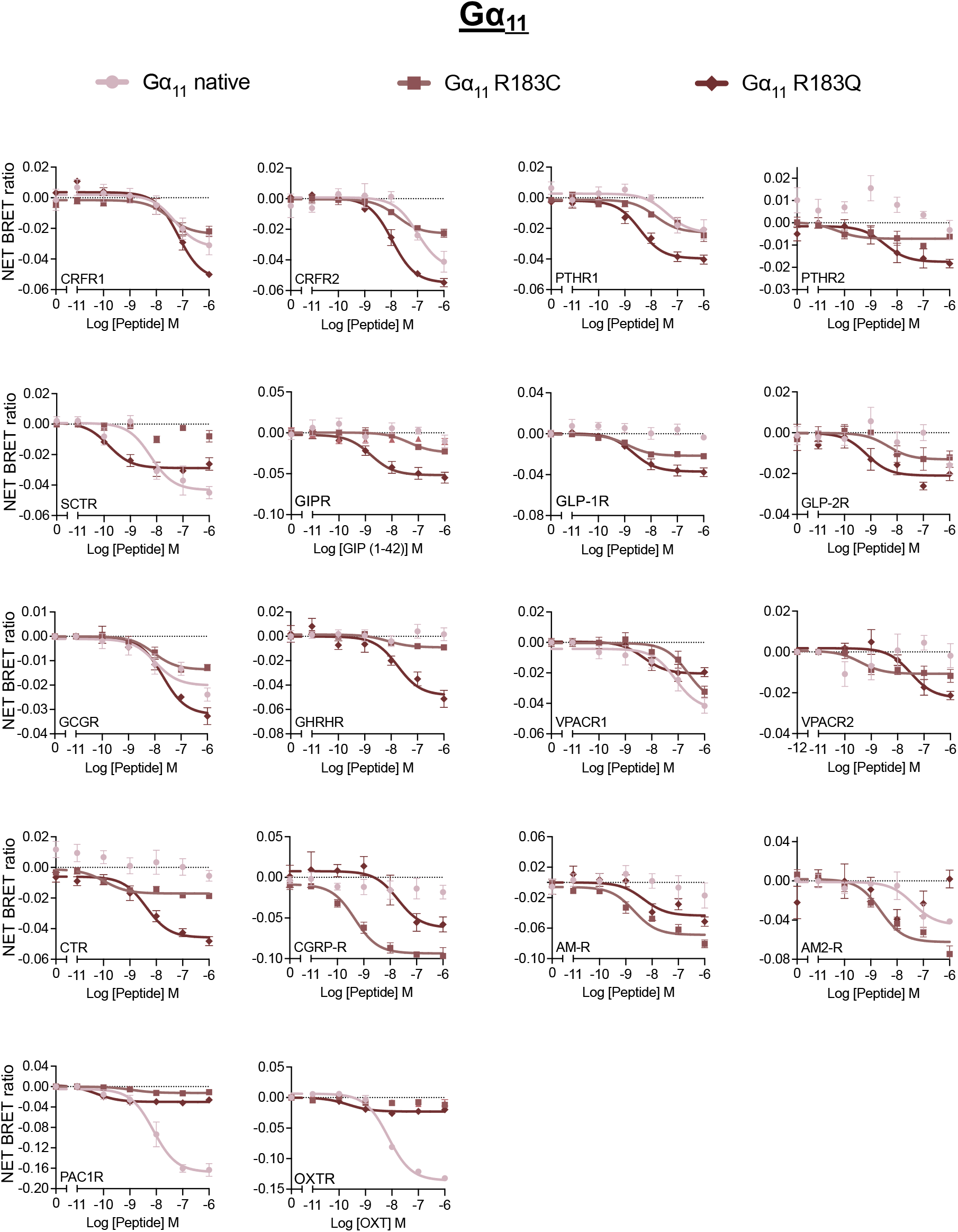
Differential responses of all class B1 GPCRs using G_11_ native and mutants in TRUPATH assay. Dose-response curves of dissociation of native or mutant G_11_ for all class B1 GPCRs. HEK 293T cells transiently transfected with each class B1 GPCR and native, R183C or R183Q G_11_ TRUPATH biosensors were stimulated with their cognate agonist. Data are the mean ± SEM of 3-6 individual repeats performed in duplicate.

**Figure 5.**
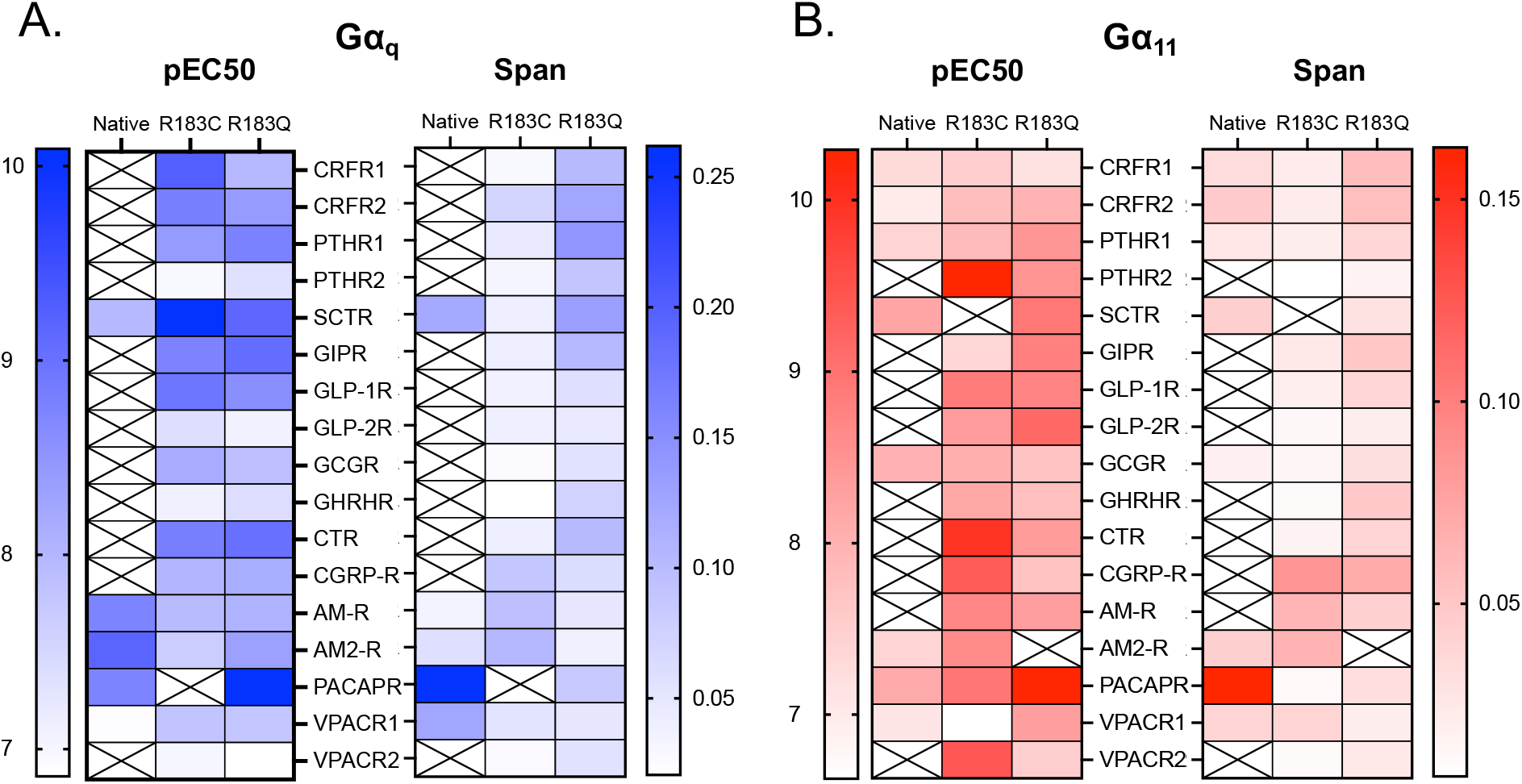
G_q/11_R183C or G_q/11_R183Q variants enhance the dynamic range for most class B1 GPCRs. pEC_50_ (left) and span (right) heatmaps of G_q/11_ native, R183C and R183Q dissociation for each class B1 GPCR in response to their cognate ligands. Data are the mean ± SEM of 3-6 individual repeats performed in duplicate.

Only secretin receptor (SCTR), vasoactive intestinal polypeptide (VPAC) receptor 1 (VPAC1R), AMR, AMR2 and pituitary adenylate cyclase-activating polypeptide receptor (PAC1R) displayed a significant dose-dependent reduction in net BRET ratio using native G_q_ (Figure 3,5). For all remaining receptors, activation of G_q_ could only be detected using the R183C and R183Q mutants. With the exceptions of glucagon-like peptide 2 receptor (GLP-2R) and CGRPR, the Gln mutant showed the greatest response. For those receptors where native G_q_ was active, the response was greatest with R183C for AMR and AMR2, and with R183Q for SCTR. Interestingly, G_q_ activation was greatest for the native protein at the PAC1R, with Gln or Cys substitution dramatically reducing the E_max_, but increasing the potency. It is of note that the PAC1R maximal response is >2-fold greater than any other receptor, suggesting that PAC1R strongly activates G_q_. It is therefore likely that, analogous to OXTR, the native response is greater than the maximal dynamic range of the mutant G_q_ proteins, bought about by their increased activity reducing the activatable pool of G_q_.

In the case of G_11_, far more class B1 GPCRs activated the native protein. Aside from GIPR and GLP-1R, native G_11_ activation was not detected for parathyroid hormone receptor 2 (PTHR2), GLP-2R, growth hormone releasing hormone receptor (GHRHR), VPAC2R, calcitonin receptor (CTR), CGRPR and AMR. When assessing the effect of the Gln and Cys substitutions, there was a very similar pattern of effects to G_q_. The use of R183Q resulted in the greatest responses for all receptors, besides the CLR-RAMP heteromers, which had the largest E_max_ for R183C; and PAC1R, where mutation of Arg183 substantially reduced the maximal response (Figure 4, 5).

It is clear from the results of this study that for GPCRs that couple strongly to G_q/11_ and for which there is significant ligand-induced loss of BRET for the native (R183) G protein, such as PAC1R, substitution of Arg is detrimental to the maximal response observed. However, for the majority of class B1 GPCRs, mutation of R183 to Cys or Gln is necessary for robust detection of G_q/11_ activation, with a receptor-specific tendency to maximally activate one or the other mutant. As summarised in Table 1, we provide guidance for the optimal G_q/11_ biosensor for each class B1 GPCR. The amino acid at position 183 that provides the greatest signalling window appears to cluster between subfamilies. For most class B1 GPCRs, Gln substitution provides the greatest signalling window. However, for the CLR-based receptors Cys substitution is most beneficial and the native Arg is optimal for PAC1R. G_q/11_ activation by the VPACRs appears minimally affected by mutation, whilst Cys and Gln substitutions are similarly beneficial for the GLP subfamily at G_q_.

**Table 1.** The summary of mutant activity as a tool to investigate Gα_q/11_ coupling to class B GPCRs

| Receptor | $G_{\alpha_q}$ | | | $G_{\alpha_{11}}$ | | |
| --- | --- | --- | --- | --- | --- | --- |
|  | native | R183C | R183Q | native | R183C | R183Q |
| CRFR1 | - | + | ++ | + | + | ++ |
| CRFR2 | - | + | ++ | + | + | ++ |
| PTHR1 | - | + | ++ | + | + | ++ |
| PTHR2 | - | - | ++ | - | - | ++ |
| SCTR | + | + | ++ | + | - | ++ |
| GIPR | - | + | ++ | - | + | ++ |
| GLP-1R | - | ++ | ++ | - | + | ++ |
| GLP-2R | - | ++ | ++ | - | + | ++ |
| GCGR | - | + | ++ | + | + | ++ |
| GHRHR | - | + | ++ | - | + | ++ |
| VPACR1 | ++ | + | + | ++ | ++ | ++ |
| VPACR2 | - | + | ++ | - | + | ++ |
| CTR | - | + | ++ | - | + | ++ |
| CGRP-R | - | ++ | + | - | ++ | + |
| AM-R | + | ++ | + | - | ++ | + |
| AM2-R | + | ++ | + | + | ++ | - |
| PAC1R | ++ | - | + | ++ | + | + |

The criteria were determined by comparing the efficacy between native and each mutant in corresponding to their cognate ligands.
(-) – no response detected
(+) – response
(++) – the optimum (largest) response detected

### R183 mutation also enables detection of G_q/11_ activation for G_s_ coupled class A GPCRs

Having demonstrated the utility of mutating R183 in G_q/11_ for class B1 GPCRs, we next wanted to investigate whether their use could be extended to class A GPCRs that have secondary G_q/11_ coupling. A recent study using BRET biosensors that detect G protein association with GPCRs, found promiscuous coupling of G_s_-coupled receptors to G_i/o_, G_q/11_ and G_12/13_ family G proteins (Okashah et al., 2019), which has also been reported in other previous studies (Kooistra et al., 2021; Okashah et al., 2019; Pándy-Szekeres et al., 2022). However, in the original TRUPATH paper (Olsen et al., 2020), no G_q_ activation was detected upon receptor activation for a selection of G_s_-coupled class A GPCRs, including the β_2_-AR that was investigated in the Okashah et al study. We, therefore, sought to investigate whether the our newly characterised mutants could resolve this discrepancy. We detected no significant loss of net BRET signal upon β_2_-AR activation in the presence of the native G_q/11_ TRUPATH biosensors (Figure 6). For both G_q_ and G_11_, introduction of either Cys or Gln resulted in a detectable, dose-dependent loss of net BRET following isoprenaline treatment. Similar to GIPR and GLP-1R, the maximal response was greater with the Q substitution than C substitution and there was a small, low potency response with G_q_ R183K. (Figure 6A, B).

**Figure 6.**
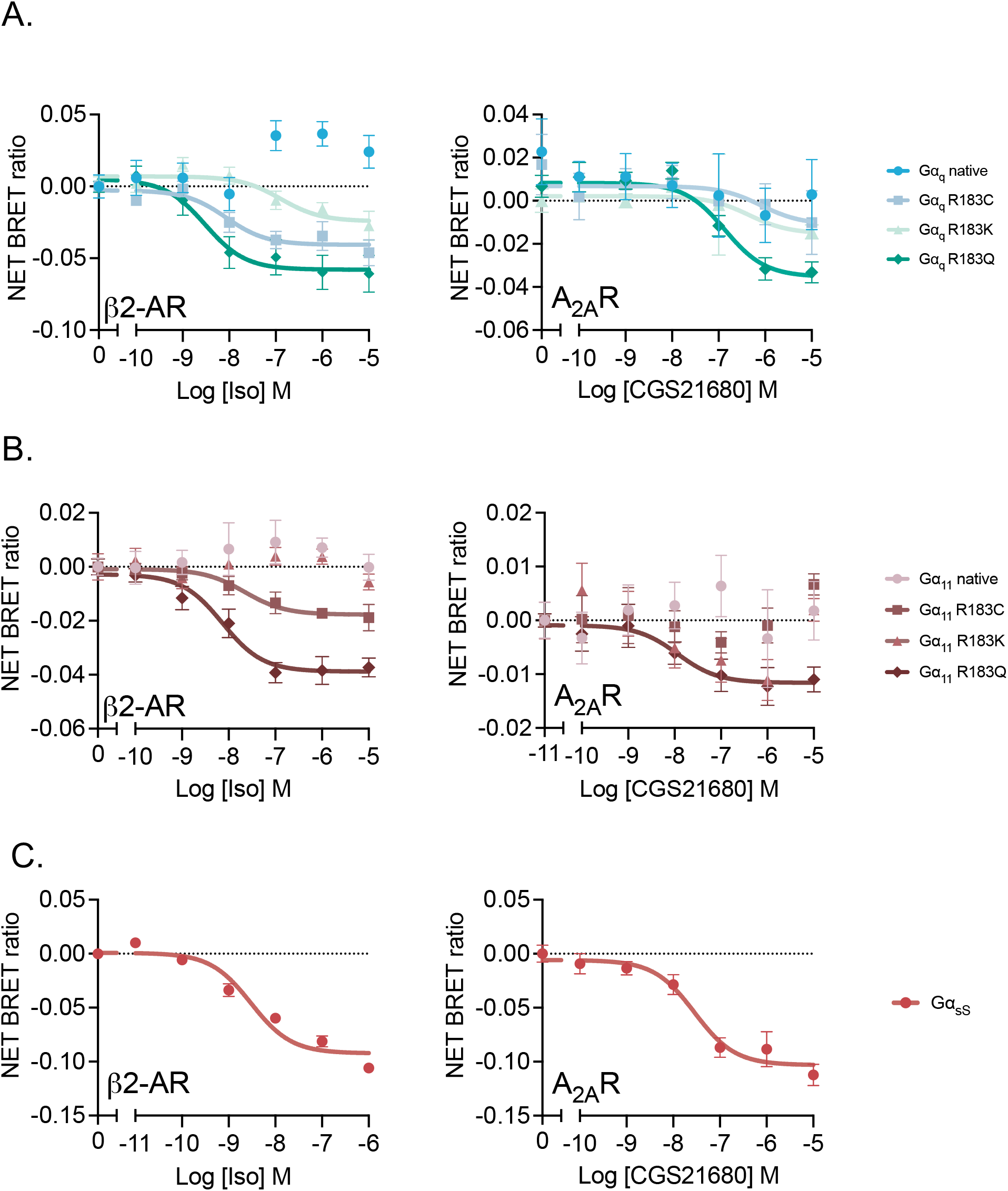
Detection of mutant G_q/11_ activation for class A G_s_-coupled receptors. (**A**-**B**) G protein dissociation in HEK 293T cells transiently expressing native, R183C, R183Q or R183K G_q_ (**A**) or G_11_ (**B**) or native G_sS_ (**C**) TRUPATH biosensors and β_2_AR or A_2A_R in response to isoprenaline or CGS21680, respectively. Data are the mean ± SEM of 3 individual repeats performed in duplicate.

We next wanted to investigate whether this was also the case for another G_s_-coupled class A GPCR that has been reported to display promiscuous secondary G protein coupling. The adenosine A_2A_ receptor (A_2A_R), is classically considered to couple to G_s_, but has been demonstrated to activate G_q_ (Inoue et al., 2019; Kooistra et al., 2021; Pándy-Szekeres et al., 2022). Similar to β_2_-AR, stimulation of either native G_q_ or G_11_ with CGS21680 (an A_2A_R-selective agonist) resulted in no significant loss of net BRET (Figure 6A-B, Native). Introduction of R183 mutations in G_q_ enabled detection of a reduction in net BRET upon agonist stimulation of A_2A_R. Dissociationthe effect of the Cys mutant was similar to that of the Lys mutant on the G_q_ subunit, with mutation to Gln showing the greatest response (Figure 6A). Interestingly, only stimulation of the R183Q of G_11_mutant resulted in a robust dose-response curve, suggesting weaker coupling of A_2A_R to G_11_ over G_q_ (Figure 6B). As a validation for normal receptor function, both β_2_-AR and A_2A_R, were assessed for G_sS_ dissociation in response to their respective agonists. As expected, a significant dose-dependent loss of net BRET was detected (Figure 6C). Overall, this indicates that the native TRUPATH system may also not be sensitive enough to detect G_q/11_ activation by class A GPCRs that primarily couple to G_s_ and thus R183 mutants can have applications beyond class B1.

To conclude, whilst downstream signalling and chemical inhibition provides evidence for G_q/11_ coupling, the use of specific mutants of Gα_q/11_ will enable further exploration of G_q/11_ signalling at class B1 GPCRs and can be expanded to corroborate the signalling profile of other GPCRs that pleiotropically couple to many G proteins.

## DISCUSSION

In the past decade, numerous biosensors have been established as probes to investigate the primary event following GPCR activation – heterotrimeric G protein interaction and dissociation (Audet et al., 2008; Avet et al., 2022; Galés et al., 2005; Inoue et al., 2012, 2019; Maziarz et al., 2020; Nobles et al., 2005; Olsen et al., 2020; Philip et al., 2007). A recent biosensor platform, TRUPATH, was developed based on the loss of BRET between Gα-Rluc8 and Gβ-GɣGFP2 upon G protein activation (Olsen et al., 2020). It has been proven that the tool is powerful to explore GPCR signalling at the G protein level (Olsen et al., 2020). However, for two typical class B1 GPCRs, GIPR and GLP-1R, which have previously been shown, and further validated here, to activate signalling pathways typical of G_q/11_ activation (Ca^2+^)_i_ mobilisation, IP1 production and NFAT-dependent gene expression, in a YM-254890-sensitive manner, no G_q/11_ activation could be detected. We demonstrate that mutation of a highly conserved residue critical to the GTPase activity of G_q/11_ (R183) increases the dynamic signalling range for class B1 GPCRs.

The G_q/11_ biosensors developed in the original TRUPATH study were validated against the promiscuous class A GPCR neurotensin-1 receptor (NTR1) (Olsen et al., 2020). Here we validated the G_q/11_ biosensors against the OXTR, a class A GPCR that primarily couples to G_q/11_ (Bichet et al., 2019; Passoni, Leonzino, Gigliucci, Chini, & Busnelli, 2016). Despite comparable (Ca^2+^)_i_, IP1 and NFAT gene expression responses from GIPR and GLP-1R, the G_q/11_ TRUPATH biosensors failed to detect a response to their cognate agonists. The fact that the OXTR, GIPR and GLP-1R responses were equally susceptible to G_q/11_ inhibition with YM-254890, suggested that the TRUPATH G_q/11_ biosensors were not sensitive enough to detect G_q/11_ responses at class B1 GPCRs that primarily couple to G_s_. Indeed, investigation of all 17 class B1 GPCRs (including CLR-RAMP complexes) revealed that <30% and <50% of receptors activated G_q_ or G_11_, respectively, despite the fact that there is evidence of G_q/11_ signalling for all class B1 GPCRs (Wootten et al., 2017). Interestingly, except for PAC1R, all responses to native G_q/11_ were relatively small compared to other TRUPATH biosensors and class A GPCR-mediated G_q/11_ activation.

There are several possible explanations for why this might occur. Firstly, it is generally considered that upon G protein activation Gα-GTP and Gβɣ subunits dissociate (Gilman, 1987), it is possible that instead there is a subtle rearrangement in conformation of Gα-GTP and Gβɣ to expose the effector interaction site that does not lead to a discernible change in the relative positions of the two tags. The latter may explain, the weak or undetectable G_q/11_ responses for most class B1 GPCRs and receptors that mainly couple to G_s_.

Within the Gα_q/11_ family, there are 3 different switches (I, II, III) that are reported to be responsible for GTP hydrolysis, with these areas becoming more rigid in the GTP-bound active conformation (Kamato et al., 2015). The presence of Rluc8 flanking these regions within Gα_q_ or Gα_11_ may impact the affinity or productivity of the interaction of G_q/11_ with GPCRs, particularly those that may bind with lower affinity to Gα_q_ than their primary G protein partner. Therefore, it is possible that addition of the BRET molecules within the Gα structure may further reduce weak G_q/11_ coupling from class B1 GPCRs that primarily couple to G_s_.

A third, most likely explanation, relates to the kinetics of G protein activation and inactivation that results in suboptimal conditions for the detection of dissociation of Gα_q/11_-Rluc8 and Gβ-GɣGFP2. Class A and class B1 GPCRs have been shown to possess distinct modes of activation upon agonist binding (Hilger et al., 2020). Whilst class A GPCRs undergo agonist-induced bending and outward movement of TM6, in class B1 GPCRs there is a breaking of the TM6 helix to create a sharp kink. This suggests that there is a higher energy barrier to activate class B1 GPCRs. Indeed, complete outward movement of TM6 in GCGR is only observed following engagement of the Gα_s_-α5 helix. Furthermore, class B1 GPCRs have been shown to possess lower guanine nucleotide exchange factor (GEF) activity compared to class A GPCRs with slower GDP dissociation and GTP binding. This is likely a result of weaker engagement of the ICL2 region with a hydrophobic pocket formed by Gα_s_ αN-β1 loop, β2-β3 loop and the α5 helix (Hilger et al., 2020). There are currently no available structures of GPCRs bound to G_q/11_. However, if the common structural changes observed upon activation and the slower GEF activity of G_s_-coupled class B1 GPCRs are similar for G_q/11_-bound receptors, it is likely that the kinetics of G_q/11_ activation by class B1 GPCRs are slower than class A GPCRs. On top of this, it has long been known that measurement of GTPase activity of native Gα_q_ is challenging, as it occurs more rapidly compared to other Gα subunits, especially under physiological conditions (Mizuno & Itoh, 2009; Mukhopadhyay & Ross, 1999). The intrinsic GTPase activity of Gα_q_ is enhanced by expression of GTPase activating proteins (GAPs), namely PLC-β, RGS4 and GAIP (Anger, Zhang, & Mende, 2004; Hepler, Berman, Gilman, & Kozasa, 1997; Kankanamge et al., 2021). The high rate of intrinsic GTPase activity coupled with endogenous expression of GAPs in HEK293T cells suggests that the kinetics of inactivation of G_q/11_ may be very high. It is therefore likely that the combination of secondary coupling of class B1 GPCRs to G_q/11_, with latent binding cavities that are potentially not optimized for efficient G_q/11_ engagement, a higher energy barrier to activation and slow activation kinetics compared to class A GPCRs and, finally, the fast inactivation kinetics of Gα_q/11_ means that the kinetic balance produces a very small dynamic range for detection of a response. This is supported by our observation that by mutating R183 in the switch I region of Gα_q/11_ to prevent intrinsic GTP hydrolysis the sensitivity of the TRUPATH system to class B1-mediated G_q/11_ activation is enhanced.

In this study, we have analysed receptor coupling to Gα_q/11_ proteins with different single residue subtitutions at position 183. These included cancer-associated mutations (Gln and Cys) and a residue (Lys) that has similar physicochemical properties compared to the native Arginine, which is conserved in all Gα subtypes and located in the switch I domain. While Lys is used to provide positively charge residue similar to Arg, the Gln mutation is more commonly found in Gα_q_, whereas the Cys mutation is more commonly associated with Gα_11_ (Murali et al., 2012; Shirley et al., 2013). Cys and Gln at position 183 have been reported to be oncogenic (Raamsdonk et al., 2010; Van Raamsdonk et al., 2009). A positive charge is known to be required to stabilise the ɣ-phosphate in the transition state of GTP hydrolysis (Wittinghofer & Vetter, 2011) and therefore, mutation of Arginine 183 has been shown to result in GTPase deficient activities in a number of Gα subtypes (Berman, Wilkie, & Gilman, 1996; Der, Finkel, & Cooper, 1986; Landis et al., 1989; Maziarz et al., 2018). Indeed, it is not surprising that mutation of this residue inhibits GTPase activity given the fact that this Arg in G_s_ is the site of ADP ribosylation by cholera toxin (Milligan & Mitchell, 1993; Palazzo, Mikolčević, Mikoč, & Ahel, 2019). Despite the loss of intrinsic GTPase activity, it has been reported that GAP activity of RGS4 is, at least partially, retained when Gα_q_ is mutated (Bastin et al., 2012; Berman et al., 1996). The results of the current study show that, for the majority of class B1 GPCRs, mutation of R183 to either Cys or Gln sufficiently changes the kinetic balance to enable detection of G_q/11_ activation. Importantly, the absence of any response for native or mutant Gα_q/11_ for three receptors known not to stimulate G_q/11_ signalling pathways (β_3_AR, D_3_R and μ-OR) (Hauser et al., 2022) provides confidence that the detected responses are not an artifact.

Interestingly, mutation of R183 in G_q/11_ reduced the range of response for OXTR but increased oxytocin potency. OXTR is primarily coupled to G_q/11_, optimally engaging G_q/11_ upon activation and, being a class A GPCR, the kinetics of G protein activation are likely faster than in class B1 GPCRs. Together, this suggests that for OXTR (and likely other G_q_-coupled class A GPCRs) the rate and extent of G_q/11_ activation is sufficient to overcome the fast inactivation kinetics to produce a detectable loss of net BRET with the native Gα_q/11_. For GPCRs that weakly activate G_q/11_, mutation of R183 to Cys or Gln increases the sensitivity of the system. Whilst this is also likely true for OXTR, the loss of heterotrimeric activatable Gαβɣ reduces the maximal signal. In effect, for strong or primary couplers of G_q/11_, mutation of R183 removes any G protein reserve, analogous to the spare receptor concept, to limit the maximal response observed. This also explains the observed increase in potency with Cys or Gln substitution from hitting the ceiling of the assay system. We, therefore, recommend that the native G_q/11_ is used for any GPCR with a robust native G_q/11_ signal. Interestingly, for two G_s_-coupled class A GPCRs that have previously been reported to promiscuously couple to G_q/11_ but show no G_q_ activation in the native TRUPATH system (Olsen et al., 2020), R183 mutation to Cys or Gln enabled detection of G_q/11_ activation, analogous to class B1 GPCRs. Thus, the fast inactivation kinetics of G_q/11_ are sufficient to reduce the signalling window for other, non-class B1, G_s_-coupled receptors that do not demonstrate primary G_q/11_ coupling. Therefore, mutation of the G_q/11_ TRUPATH biosensors appears necessary for robust, reliable detection of G_q/11_ activation of G_s_-coupled receptors across different families of GPCRs.

As previously discussed, mutation of R183 allowed enhanced detection of G_q/11_ activation for most class B1 GPCRs. However, this was not the case for PAC1R, which showed a reduced range and increased potency of response with the mutants. The large and potent native response coupled to the similar effects of R183 mutation for PAC1R to OXTR indicates that PAC1R strongly couples to G_q/11_ and engages it with high affinity upon agonist-induced activation. The native response is thus greater than the system maximum with the mutants. Alignment for PAC1R shows differences in the ICL2 regions, particularly in comparison with GIPR. The length of ICL2 has been reported to be a key determinant and important contact site between G proteins and the receptor upon activation (Liang et al., 2020). These differences indicate that differential conformational changes in mediating G protein engagement may occur within the same GPCR class. However, to validate this hypothesis, further experiments would be required. For those receptors with enhanced net BRET signal with Arg substitutions, the dynamic range was generally greatest for Gln for both G_q_ and G_11_. Interestingly there were some differences between subfamilies; CLR-based receptor complexes displayed the greatest range with Cys substitution; GLP1/2R showed minimal improvement with Gln over Cys for G_q_. These differences are most likely due to subtle differences in the conformation of the receptor when bound to G_q/11_ and the strength of interaction sites. Despite the possibility that clustering effect may be dependent on their phylogenetic tree, there is currently no evidence to explain preferential coupling amongst these receptors (Supplementary Figure S6). However, our results indicate that the occurrence of these mutations in different cancer types may not result in a generalised increase of G_q/11_ coupling across receptors but could elicit receptor-specific signalling alterations.

In summary, our approach reveals that a modification to the newly developed TRUPATH system allows detection of reliable and robust G_q/11_ responses for all class B1 GPCRs and has utility with other, non-class B1, G_s_-coupled GPCRs. Whilst native Gα_q/11_ is sufficient for VPAC1R and PAC1R, the dynamic range is greater for CLR-based receptors using the R183C mutant, and most other class B1 GPCRs are optimal with R183Q. Given the fact that other studies have also failed to detect any G_q/11_ activation for GIPR, GLP-1R or CRFR1 (Avet et al., 2022; Jones et al., 2021), it is possible that mutation of R183, to reduce intrinsic GTPase activity and generate a more stable Gα-GTP may enable reliable detection of G_s_-coupled GPCR-mediated G_q/11_ activation.

## AUTHOR CONTRIBUTIONS

GL, DS and MH conceived and designed the research; DS, MH, AP, XH, MR, EW, KB and MDF performed the experiments, DS, MH and GL wrote the manuscript, all authors revised, edited and approved the manuscript.

## ACKNOWLEDGEMENT

We would like to thank the following sources of funding. The Leverhulme Trust (Grant RPG-2017-255 to fund KB), AstraZeneca for funding MR and EW, while AP is funded by an (BBSRC)-iCase studentships (AP: BB/ JO14540/1) with AstraZeneca, and a China Scholarship Council Cambridge International Scholarship for XH. We thank Takeda Development Centre Americas Inc for funding DS. MMS acknowledges support from the Wellcome Trust Institutional Strategic Support Fund and the Isaac Newton Trust [22.23(d)]. GL is funded by a Royal Society Industry Fellowship.

