## Supporting Info for "Cancer-Associated Mutations Enhance The Sensitivity Of The Trupath Gα_Q/11_ System"

Supplementary Figures 6.

Figure S1.

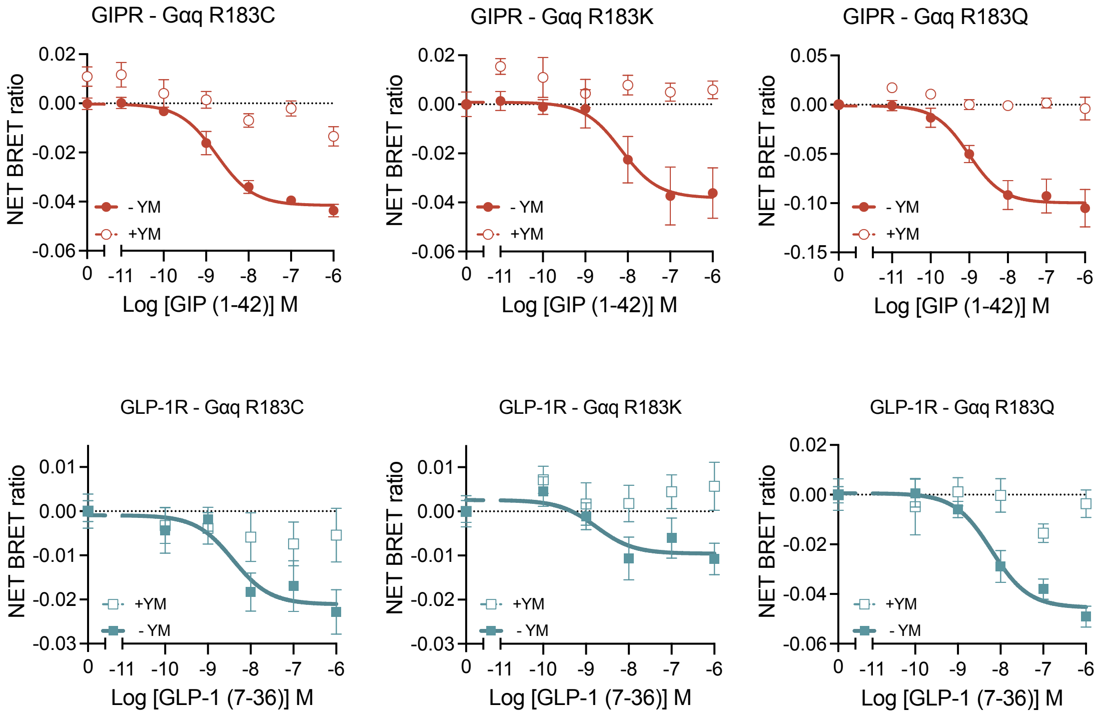


**Supplementary Figure 1: YM254890 inhibits coupling of $G_q$ variants to GIPR and GLP-1R in TRUPATH assay**

$G_{q/11}$ mutants are sensitive to YM254890 inhibition. HEK293T cells transiently transfected with GIPR or GLP-1R and the native, R183C, R183K and R183Q $G_{q/11}$ TRUPATH biosensors were stimulated with GIP (1-42) or GLP-1 (7-36)$NH_2$ in the absence or presence of 100 nM YM-254890. Data are the mean ± SEM of 3 individual repeats performed in duplicate.

Figure S2

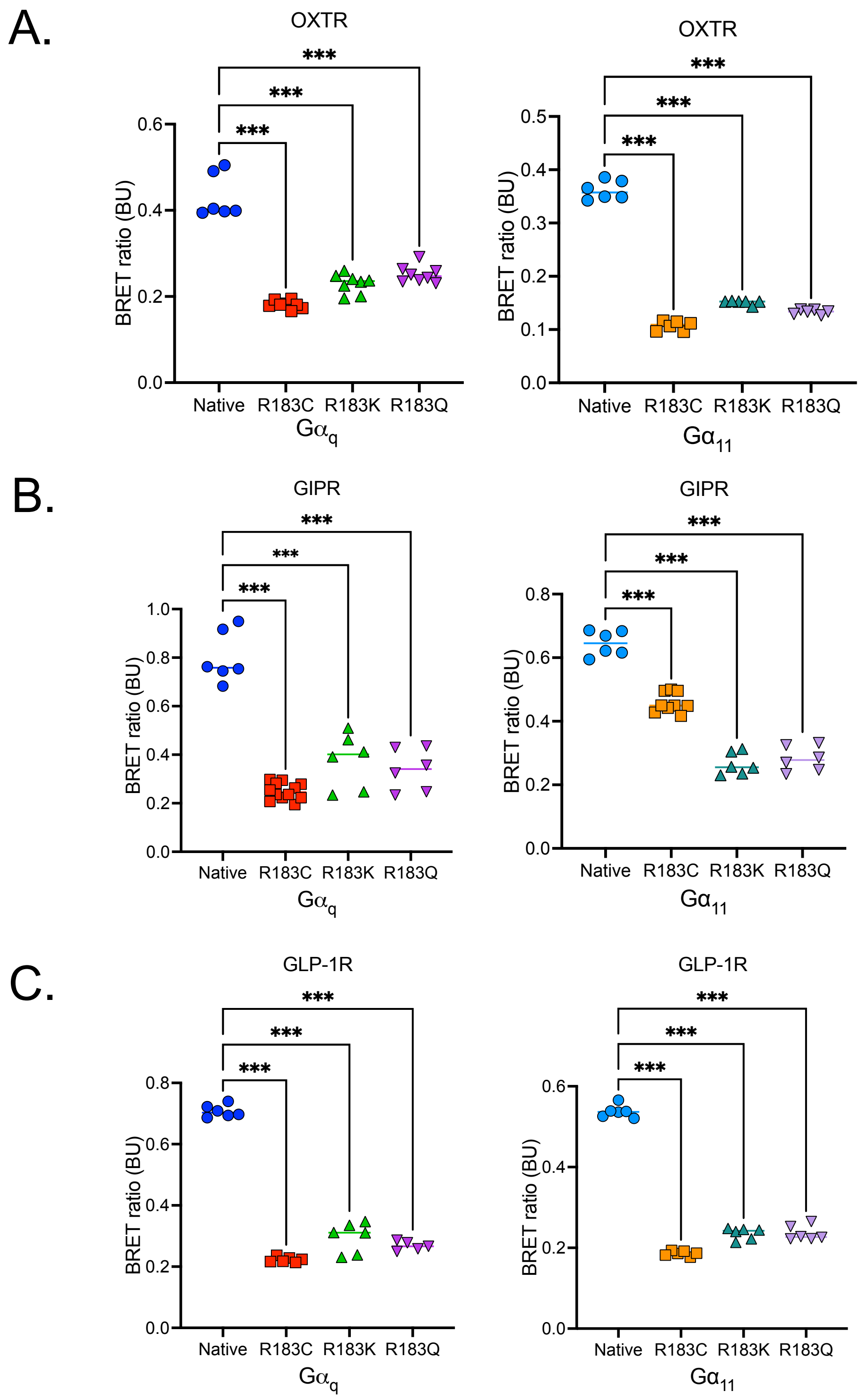


**Supplementary Figure 2: Mutation of R183 reduces the basal BRET ratio**

Raw BRET values for native, R183C, R183K and R183Q $G_{q/11}$ TRUPATH biosensors in the absence of agonist (vehicle treated) from OXTR (A), GIPR (B), and GLP-1R (C), which are displayed in Figure 2. Data are the mean ± SEM of 3-6 individual repeats performed in duplicate.

Figure S3.

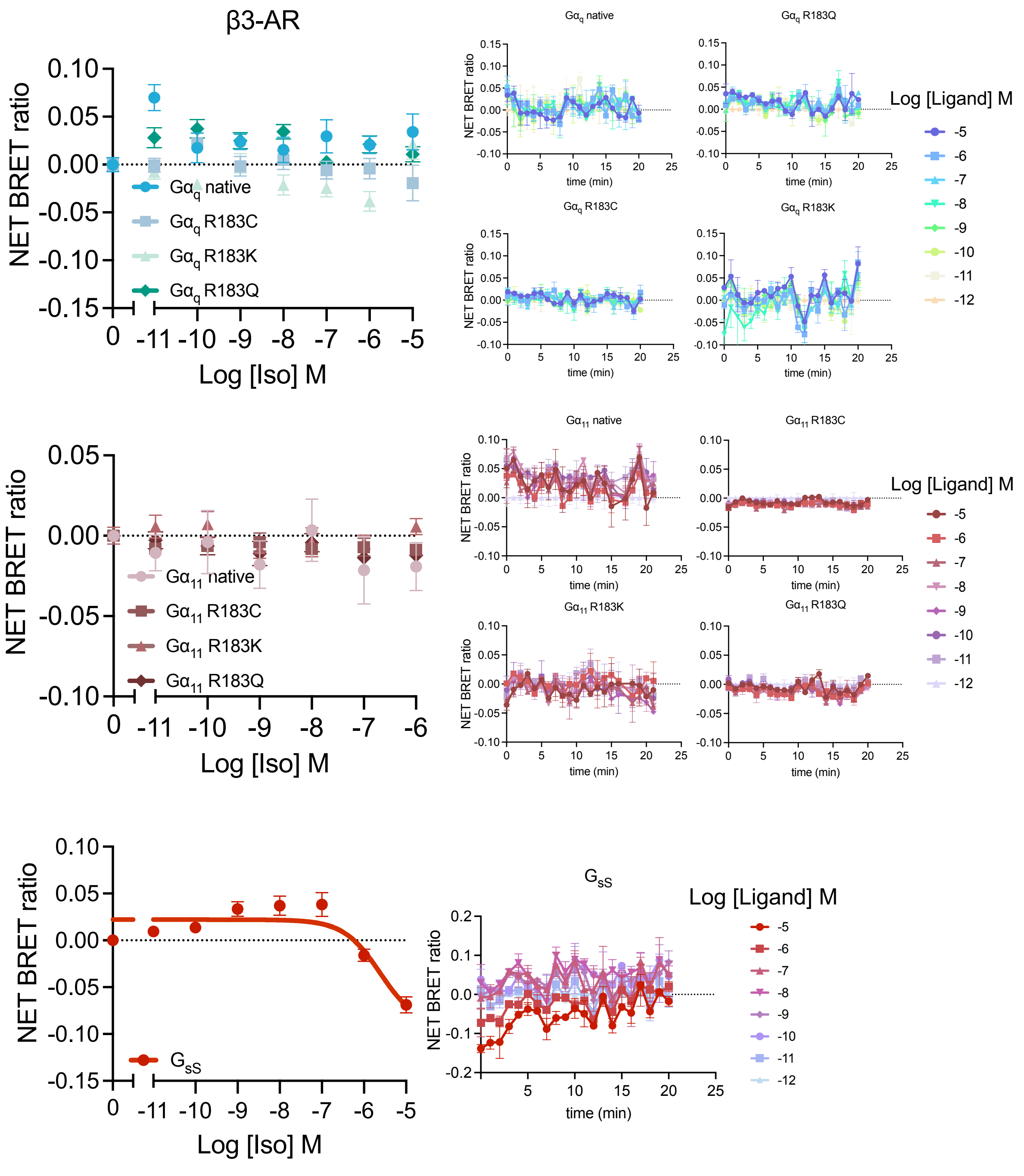


**Supplementary Figure 3: Curves and traces from receptors that are not coupled to $G_q$/$G_{11}$ - $\beta_3$AR**

Activation of $\beta_3$-AR does not stimulate dissociation of native or mutant $G_{q/11}$. (**A**) Dose-response curves (left) and temporal traces (right) of isoprenaline stimulated native and mutant $G_{q/11}$ TRUPATH biosensor dissociation in HEK293T cells transiently transfected with $\beta_3$-AR. (**B**) HEK293T cells transiently transfected with $\beta_3$-AR and $G_{sS}$ TRUPATH biosensor show rapid, dose-dependent loss of net BRET in response to isoprenaline. (**C**) Absence of isoprenaline stimulated intracellular calcium mobilisation in HEK293T cells transiently expressing $\beta_3$-AR. Data are the mean ± SEM of 3-6 individual repeats performed in duplicate.

Figure S4.

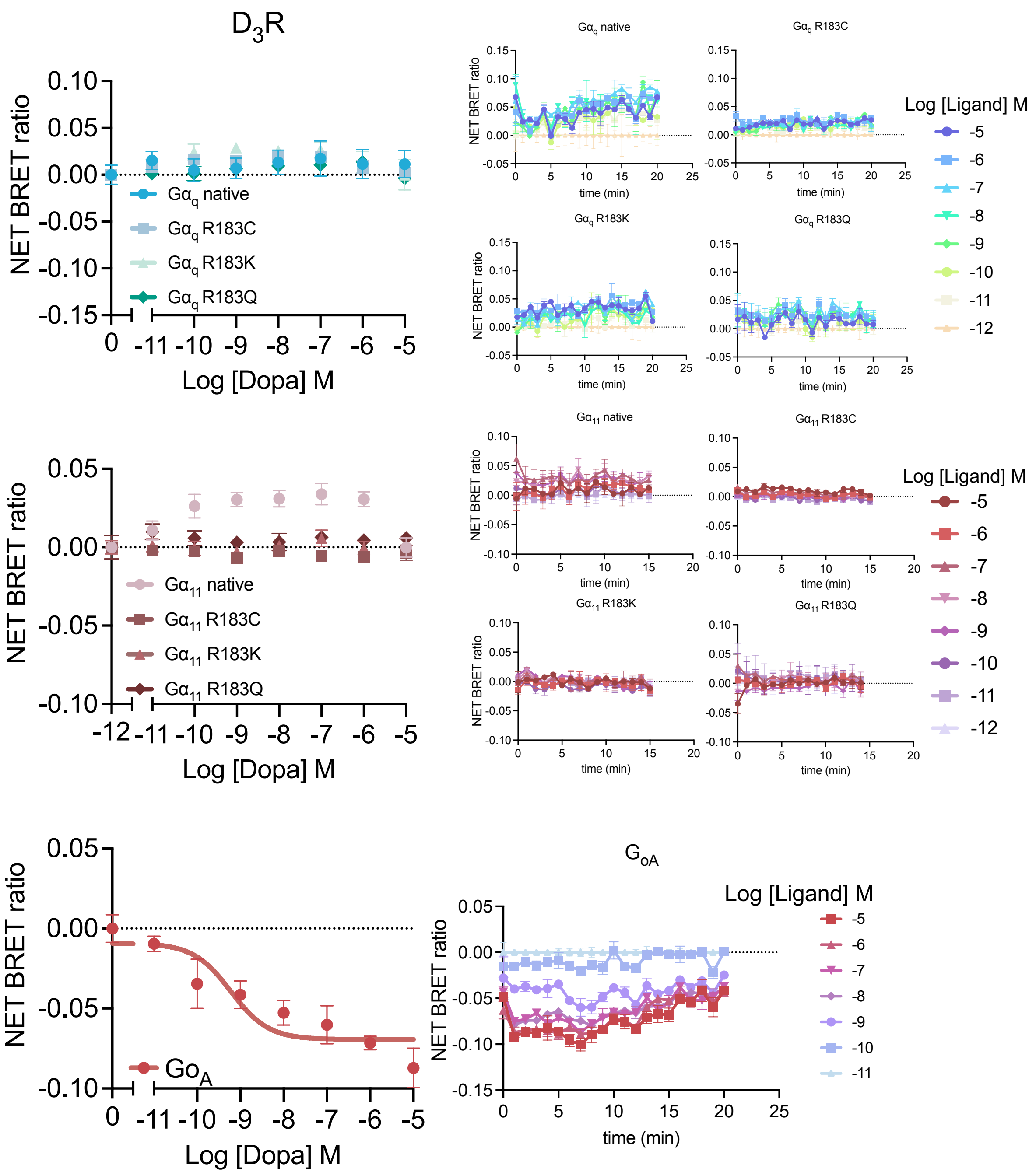


**Supplementary Figure 4: Curves and traces from receptors that are not coupled to $G_q/G_{11}$ – $D_3R$**

Activation of $D_3R$ does not stimulate dissociation of native or mutant $G_{q/11}$. (**A**) Dose-response curves (left) and temporal traces (right) of dopamine stimulated native and mutant $G_{q/11}$ TRUPATH biosensor dissociation in HEK293T cells transiently transfected with $D_3R$. (**B**) HEK 293T cells transiently transfected with $D_3R$ and $G_{oA}$ TRUPATH biosensor show sustained, dose-dependent loss of net BRET in response to dopamine. (**C**) Absence of dopamine stimulated intracellular calcium mobilisation in HEK293T cells transiently expressing $D_3R$. Data are the mean ± SEM of 3-6 individual repeats performed in duplicate.

Figure S5.

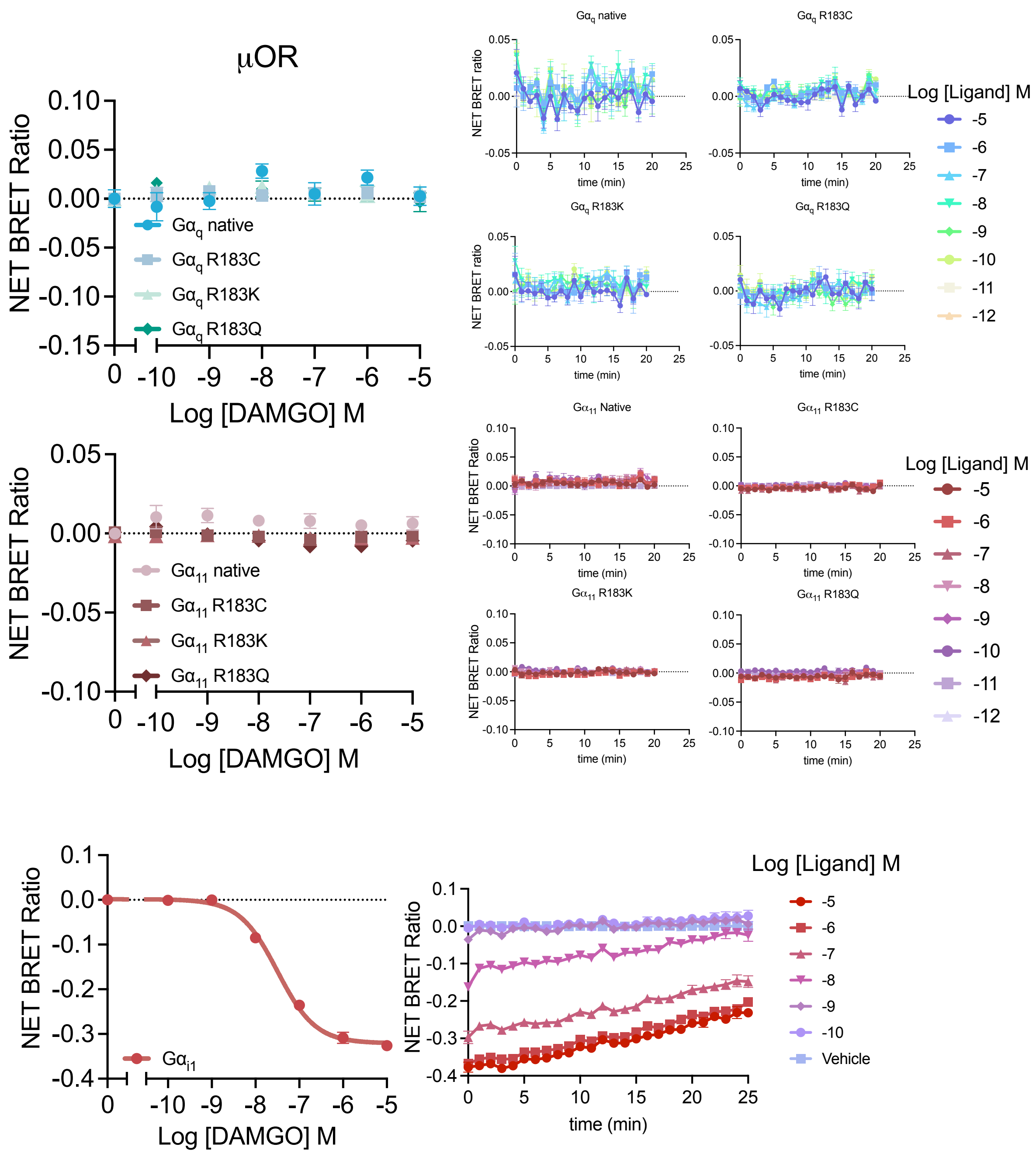


**Supplementary Figure 5: Curves and traces from receptors that are not coupled to $G_{q/11}$ –μ-OR**

Activation of μ-OR does not stimulate dissociation of native or mutant $G_{q/11}$. (**A**) Dose-response curves (left) and temporal traces (right) of DAMGO stimulated native and mutant $G_{q/11}$ TRUPATH biosensor dissociation in HEK 293T cells transiently transfected with μ-OR. (**B**) HEK 293T cells transiently transfected with μ-OR and $G_{i1}$ TRUPATH biosensor show sustained, dose-dependent loss of net BRET in response to DAMGO. (**C**) Absence of DAMGO stimulated intracellular calcium mobilisation in HEK 293T cells transiently expressing μ-OR. Data are the mean ± SEM of 3-6 individual repeats performed in duplicate.

Figure S6.

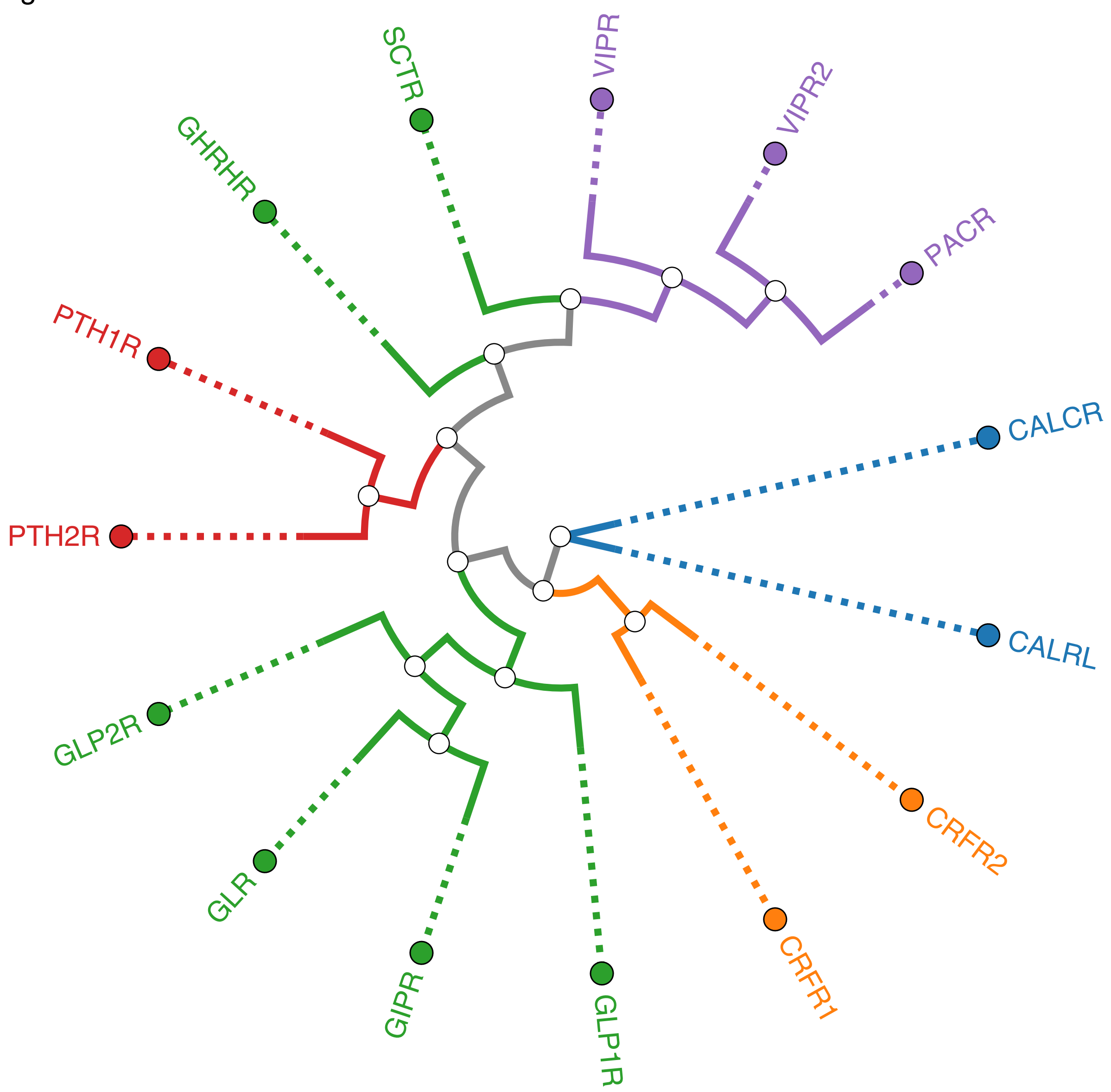


**Supplementary Figure 6: Phylogenetic tree of receptors within class B1.**

Application of $G_{q/11}$ mutants in TRUPATH assay displayed clustering effect may be related to their similarities in phylogenetic tree.
